# Genomic characterization of an atypical hydrogen sulfide-negative *Salmonella enterica* serovar Senftenberg strain lacking somatic antigen expression isolated from cooked mussels

**DOI:** 10.64898/2026.03.06.710058

**Authors:** Alexandre Lamas, Antonio Lozano-León, Alejandro Garrido-Maestu, Narjol Gonzalez-Escalona

**Affiliations:** Food Hygiene, Inspection and Control Laboratory (Lhica), Department of Analytical Chemistry, Nutrition, and Bromatology, Veterinary School, Campus Terra, Universidade de Santiago de Compostela (USC), 27002, Lugo, Spain; Health & Nutrition Lab Manager. SGS Seafood Lab. Polígono Industrial Castiñeiras, Av. Castiñeiras, 30, 36939 Bueu, Pontevedra; Laboratory of Microbiology and Technology of Marine Products (MicroTEC), Institute of Marine Research (IIM-CSIC), Vigo, Spain; Genomics Development and Applications Branch, Division of Food Safety Genomics, Office of Applied Microbiology and Technology (OAMT), Office of Laboratory Operations and Applied Science (OLOAS), Human Foods Program, Food and Drug Administration, College Park, MD USA

**Keywords:** *Salmonella enterica*, atypical strains, somatic antigens, H_2_S-negative, whole genome sequencing, hybrid assembly, mussels, food safety, One Health

## Abstract

Atypical *Salmonella enterica* strains that evade conventional detection pose significant challenges to food safety surveillance. A hydrogen sulfide (H₂S)-negative and serologically untypable S. enterica strain (SF1060) was detected by qPCR from cooked farmed mussels in Galicia, Spain, and characterized using phenotypic and genomic approaches. Despite typical biochemical profiles, SF1060 failed to produce black colonies on XLD agar and lacked detectable somatic antigens by conventional serotyping. Hybrid genome assembly using Nanopore and Illumina sequencing yielded a closed chromosome and five plasmids. *In silico* analyses identified the strain as *S.* Senftenberg ST14. Comparative genomics revealed a chromosomal inversion at the *rfb* operon (encoding enzymes needed to synthesize deoxysugars and O antigens) mediated by *IS5*-family transposase *ISEc68*, which truncated the *rfbD* gene and separated the reminding *rfb* genes at *rfb*D, disrupting O-antigen biosynthesis, explaining the serotyping failure. The *phs* operon responsible for H₂S production lacked premature stop codons, suggesting the H₂S-negative phenotype results from an alternative mechanism. This study demonstrates how whole-genome sequencing resolves identification of atypical strains that fail culture-based detection and emphasizes the critical need for molecular surveillance methods in seafood safety programs, particularly in regions where atypical *S. enterica* variants may be endemic.

**Importance Statement:** Pathogen surveillance is in a constant race against microbial evolution. The phenotypic methods used to detect and isolate foodborne bacteria like *Salmonella enterica* from foods are effective, but only if the pathogens retain their expected characteristics. This study provides a clear genetic snapshot of how *Salmonella enterica* can adapt to evade detection. This study characterizes a strain from a marine environment that had undergone genetic rearrangement, leading to the concurrent loss of a key biochemical marker and the somatic antigens essential for serological typing. Crucially, only genomic analysis could provide a definitive explanation for this serotyping failure; the underlying mechanism would have remained unknown using conventional methods alone. This work demonstrates that relying on a fixed set of phenotypic traits is an increasingly fragile strategy and highlights why genomic surveillance must become a routine component of food safety programs to keep pace with pathogen evolution.

## Introduction

*Salmonella enterica* is one of the most common pathogens responsible for foodborne illnesses worldwide. More than ∼80,000 cases are annually reported in Europe (1) , and more than 1 million in the US (2). These figures highlight the importance of making its detection, and accurate characterization, critical for food safety. Its detection in foodstuffs is based on cultured-based methods such as ISO 6579 or BAM Chapter (3, 4). Even though these methods are highly reliable, their performance is reduced when dealing with atypical strains which may not grow on selective media with the expected appearance. The implementation of molecular methods, such as real-time PCR (qPCR), which are not affected by the phenotype, are a way to overcome this limitation of classical methods.

Mussel aquaculture is a key industry in Spain, which is ranked among the top five producers worldwide, and more specifically in Galicia. This region is located on the northwest of the country, and it is responsible for more than 90 % of the production of this product (5). The marine environment presents unique challenges for *S. enterica* surveillance, as the pathogen can persist in seawater, biofilms, and filter-feeding shellfish, creating complex transmission networks that link environmental reservoirs, wildlife, and human consumers. These figures highlight the importance of assuring the safeness of the product to safeguard the consumers and avoid the potential economic impact due to potential foodborne diseases.

The presence of atypical *S. enterica* strains has already been reported in this area (6), and even strains bearing genes providing resistance to last resource antibiotics like colistin (7). This fact indicates that the implementation of phenotype independent techniques is needed in order to better control this pathogen. In this sense, methodologies based on the techniques previously mentioned, PCR/ qPCR, result of high interest, but with the advancement of novel molecular techniques, such as Whole Genome Sequencing (WGS) provides a comprehensive tool for characterizing atypical strains, offering insights into their genetic basis and improving surveillance efforts (8).

During routine testing for the presence of *S. enterica* in mussels from Galicia, Spain, a particular strain was detected that failed to produce the typical black colonies on XLD agar, despite being identified as *S. enterica* by qPCR. Subsequent analysis at a reference laboratory revealed that this strain was untypable by conventional serotyping methods. These unusual phenotypic and serological characteristics prompted a detailed genomic investigation to elucidate the genetic basis underlying the absence of hydrogen sulfide (H₂S) production and the untypable serotype. Here, we describe the genomic features of this atypical strain and discuss the possible molecular mechanisms responsible for its distinctive phenotype. The objectives of this study were to: (i) characterize the genomic features of an atypical H₂S-negative *S. enterica* strain from cooked mussels, (ii) identify the genetic mechanisms responsible for loss of somatic antigen expression and H₂S production, and (iii) assess the phylogenetic relationship of this strain to other *S.* Senftenberg strains from the region.

## Materials and Methods

### Sample collection and bacterial isolation

Samples were collected as part of routine microbiological monitoring of mussel production facilities in Galicia, Spain, during July 2019 - July 2020). A total of 1,665 samples were screened during this surveillance period. *S. enterica* strain SF1060 was isolated from routine microbiological controls of mussels. Briefly, 25 g of mussel were mixed with 225 mL of Buffered Peptone Water (BPW, BioMérieux, S.A., Marcy l’Etoile, France) and homogenized for 90 s in a stomacher, then the mixture was incubated at 37 °C for 18 ± 2 h. The DNA from the enriched samples were extracted with the PrepSEQ™ Rapid Spin Sample Preparation Kit (Applied Biosystems, Foster City, CA, USA) and then screened for *S. enterica* using the MicroSEQ™ *Salmonella enterica* detection Kit (Applied Biosystems, Foster City, CA, USA) following the protocol provided by the manufacturer in a 7500 Fast Real Time PCR System Thermal Cycler (Applied Biosystems, Foster City, CA, USA). Samples returning a positive qPCR result were confirmed by streaking the enriched sample on XLD (BioMérieux, S.A., Marcy l’Etoile, France) and Brilliance™ *Salmonella* agar (OXOID, Hampshire, UK). The plates were incubated at 37 °C overnight. Typical *S. enterica* colonies were re-isolated onto Tryptic Soy Agar (TSA, Merck KGaA., Germany) and incubated at 37 °C overnight. For *S. enterica* confirmation, biochemical profile. was performed with the API20E (BioMérieux, S.A., Marcy l’Etoile, France) and agglutination test was performed with *Salmonella* test kit (Oxoid Diagnostic Reagents, Oxoid, UK). *S. enterica* serotyping was performed using the Kauffman-White typing scheme by slide agglutination for the detection of somatic (O) and flagellar (H) antigens with standard antisera in an external accredited laboratory. *S. enterica* strain SF1060 was conserved in cryovials at -20 °C until use.

### DNA isolation and quantification

For DNA isolation, *S. enterica* strain SF1060 was streak in a Brain Heart Infusion (BHI) agar plate (Merk Millipore, Germany) and incubated 24h at 37°C. A single colony was resuspended in 10 mL of BHI broth and incubated at 37 °C for 18 h at 130 rpm. Finally, 2mL were transferred to a microtube and centrifuged at 16000 ×g for 2 min. The supernatant was discarded, and pellet used for DNA extraction with PureLink™ Genomic DNA Mini Kit (Invitrogen, Carlsbad, CA, USA) following the protocol described for Gram-negative bacteria. Qubit^™^ dsDNA Quantitation, Broad Range in combination with Qubit^™^ 4 was used for DNA quantification.

### Whole-genome sequencing

Long read sequencing was performed using Nanopore sequencing using the MinION™ Mk1C device [Oxford Nanopore Technologies (ONT), Oxford, UK]. Briefly, 400 ng of pure DNA extracted from *S. enterica* strain SF1060 were used to prepare the DNA library using the Rapid Barcoding Sequencing kit (SQK-RBK004, Oxford Nanopore Technologies (ONT)). The sample was run in a FLO-MIN106 (R9.4.1) flow cell (FAV41850), according to the manufacturer’s instructions, for 6 hours (Oxford Nanopore Technologies). The Dorado basecaller v7.1.4 was used for basecalling the run output using the High-accuracy basecalling model (HAC, basecall_model— dna_r10.4.1_e8.2_260bps_sup@v3.5.2). The reads < 4000 bp and quality scores of <9 were discarded for downstream analysis using Fitlong v0.2.1 (https://github.com/rrwick/Filtlong) with default parameters. The circular closed genome for the SF1060 strain was obtained by de novo assembly using the Flye program v2.9.2 (9), using the parameters for HAC: --nano-hq -i 4. The short-read sequencing was performed by Biomarker Technologies (BMKGENE GmbH Münster, Germany). The DNA library was prepared with VAHTS Universal DNA Library Prep Kit for Illumina V4 (Vazyme Biotech Co., Ltd., Nanjing, China) and sequenced in an Illumina NovaSeq X (Illumina, San Diego, CA, USA) with the NovaSeq X Series Reagent Kit using 2 × 150 bp paired-end chemistry according to the manufacturer’s instructions, at a 100X coverage. Reads were trimmed with Trimmomatic v0.36 (10) using default parameters.

### Genome assembly and annotation

The complete circular closed polished genome for the SF1060 strain was obtained by a *de novo* hybrid assembly using both nanopore and Illumina data for that strain with Unicycler v0.5.0 (11). Unicycler assembled the chromosome as circular closed and oriented the chromosome to start at the *dnaA* gene. The annotation of the SF1060 genome was performed using Prokka v1.14.6 (12).

## Bioinformatic analyses

### *In silico* serotyping, antimicrobial resistance, and virulence

MLST and serotyping. *S. enterica* MLST was determined using the Enterobase website (https://enterobase.warwick.ac.uk/species/index/senterica) and the serotype using Seqsero2 (13). The antimicrobial resistance genes (AMR) gene detection was performed using ResFinder v4.6.0 (http://genepi.food.dtu.dk/resfinder) (14) [14]. The presence of *S. enterica* Pathogenicity Islands (SPIs) was determined with the tool SPIFinder 2.0 (15).

### Phylogenetic analysis

The phylogenetic relationship of the strains was assessed by a whole genome multi-locus sequence typing (wgMLST) analysis using Ridom SeqSphere+ v9.0.8. The genome of *S.* Senftenberg CFSAN002050 (NC_021818.1) (4103 CDSs) was used as reference. Genes that were repeated in more than one copy in any of the two genomes were removed from the analysis as failed genes. A task template was then created that contained both core and accessory genes for each reference strain for any future testing. Each locus (core or accessory genes) from the reference strain was assigned allele number 1. The SF1060 hybrid assembly in this study was queried against the task template. If the locus was found and was different from the reference genome or any other queried genome already in the database, a new number was assigned to that locus and so on. After eliminating any loci that were missing or found having indels (showed as failed call in the software) from the genome of any strain used in our analyses, we performed the wgMLST analysis. These remaining loci were considered the core genome shared by the analyzed strains. We used Nei’s DNA distance method to calculate the matrix of genetic distance, considering only the number of same/different alleles in the core genes. A neighbor-joining (NJ) tree was built using pairwise ignoring missing values and the genetic distances after the wgMLST analysis. wgMLST uses the alleles number of each locus to determine the genetic distance and build the phylogenetic tree. The use of allele numbers reduces the influence of recombination in the dataset studied and allows for fast clustering determination of genomes. The list of the genomes used for phylogenetic analysis is listed in Supplementary Table 1.

### Comparative genomics

CLC Genomics Workbench v20 (QIAGEN, Redwood City, USA) was used for creating the genome comparison SF1060 against *S. enterica* subsp. *enterica* serovar Senftenberg strain GTA-FD-2016-MI-02533-2 (CP038604).

## Results

### Detection and phenotypic characterization of atypical *S. enterica* strain

During the sampling process for *S. enterica* detection from frozen cooked mussels, a qPCR positive result with a Cq value of 21 was obtained in one of the samples. This sample, however, when streaked into the selective *S. enterica* media, did not show the typical, black-centered colonies on XLD agar. On the other hand, when streaked unto another selective media for *S. enterica* (Brilliance™ *Salmonella*), the expected typical morphology and color (purple) was observed. A further qPCR reaction also showed a positive result for *S. enterica* These two results showed that this was indeed a *S. enterica* strain, albeit atypical, and that this cooked mussel sample was indeed positive for *S. enterica* (Figure 1). The *S. enterica* identity for this sample was further confirmed using an API20E test showing a typical biochemical profile for *S. enterica*, except for the production of H_2_S, which was negative also in this test. The isolated strain was then serotyped in an accredited laboratory using commercial antisera against somatic and flagellar antigens (Kauffman-White Scheme). Only the antigens of the first flagellar phase were detected, and it was not possible to detect somatic antigens. The final formula obtained with agglutination test was - : g, s, t : -.

**Figure 1.**
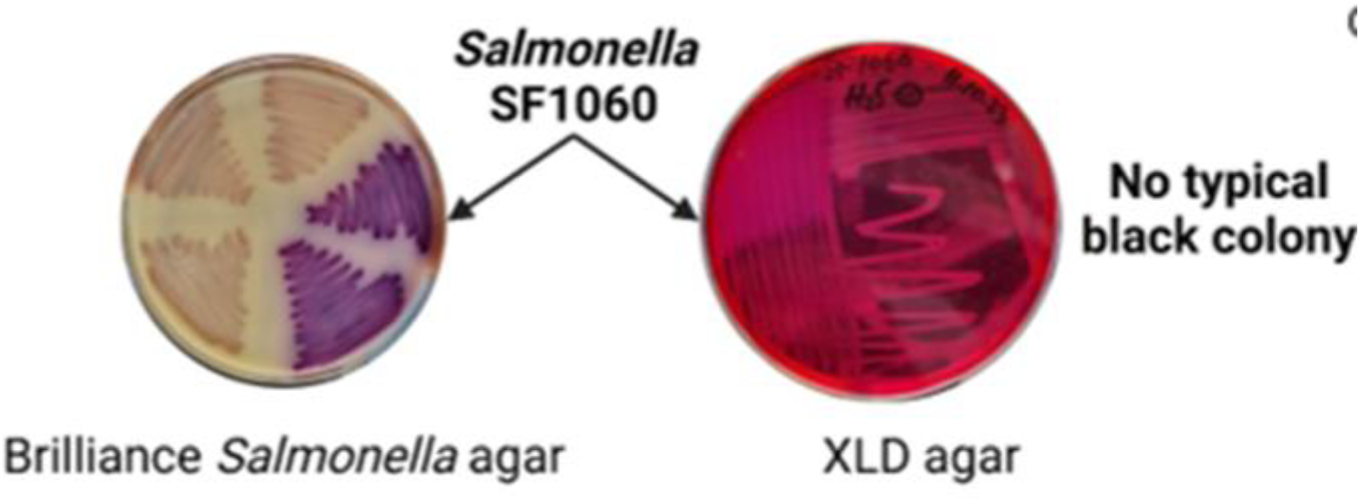
Colony morphology of *S. enterica* serovar Senftenberg strain SF1060 on selective media. (Left) Purple colonies on Brilliance™ *Salmonella* agar showing typical *S. enterica* morphology. (Right) Colonies on XLD agar lacking the characteristic black center due to absence of H₂S production, demonstrating the atypical phenotype that complicates culture-based detection.

### Genome sequencing and de novo assembly

The entire closed genome of SF1060 was obtained by whole genome sequencing and consisted of 5 contigs in total (1 chromosome and 5 plasmids) for a total length of 5,018,559 bp. The chromosome size was 4,883,048 bp with a GC% of 52.0 and annotation with Prokka showed 4834 annotated genes, 22 rRNA, 1 tmRNA and 89 tRNA and 2 CRISPR region (Table 1).

**Table 1.** Genomic characteristics of *S. enterica* serovar Senftenberg strain SF1060. The complete genome was assembled using hybrid assembly (Oxford Nanopore and Illumina sequencing) and consists of one chromosome and five plasmids. GC content (%) of each contig.

| Contig | Size | <sup>a</sup>GC% | Genes |
| --- | --- | --- | --- |
| 1 | 4,883,048 | 52.0 | 4834 |
| 2 | 71,607 | 53.2 | 82 |
| 3 | 50,564 | 53.7 | 57 |
| 4 | 7,539 | 49.0 | 10 |
| 5 | 3,506 | 51.3 | 3 |
| 6 | 2295 | 50.5 | 3 |

<sup>a</sup>Bacterial genome GC content.

### *In silico* characterization and antimicrobial resistance profiling

*In silico* serotyping and MLST analyses identified this strain as serotype Senftenberg - somatic and flagellar antigens (1,3,19:g,s,t:-) and belonging to ST14. *In silico* AMR prediction showed the presence of the aac(6’)-Iaa, a cryptic gene in *S. enterica* Also SPI-1, SPI-2, SPI-3, SPI-4, SPI-5, SPI-9 were detected.

### Genetic basis for loss of somatic antigen expression

The *rfb* gene cluster is responsible for somatic antigen biosynthesis in *S. enterica*. A Pairwise genome alignment with another Senftenberg strain GTA-FD-2016-MI-02533-2 (CP038604) showed a chromosomal re-arrangement that occurred at the *rfb* operon, responsible for O-antigen biosynthesis. The *rfb* gene cluster of strain GTA-FD-2016-MI-02533-2 (CP038604) is composed by genes, *rfbA*, *rfbB rfbC,* and *rfbD.* This re-arrangement occurred at the gene *rfbD,* which encodes a dTDP-4-dehydrorhamnose reductase (Uniprot P26392), mediated by a IS5 family transposase ISEc68, resulting in a truncated *rfbD* gene (Figure 2). Thus, the protein of the resulting CP038604 genome would have 300 residues while the protein codified by SF1060 would have 297 amino acids according to the Prokka annotation. This probably renders the gene inactive and without the expression of that protein resulting in the loss of the somatic antigens in the bacterial surface.

**Figure 2.**
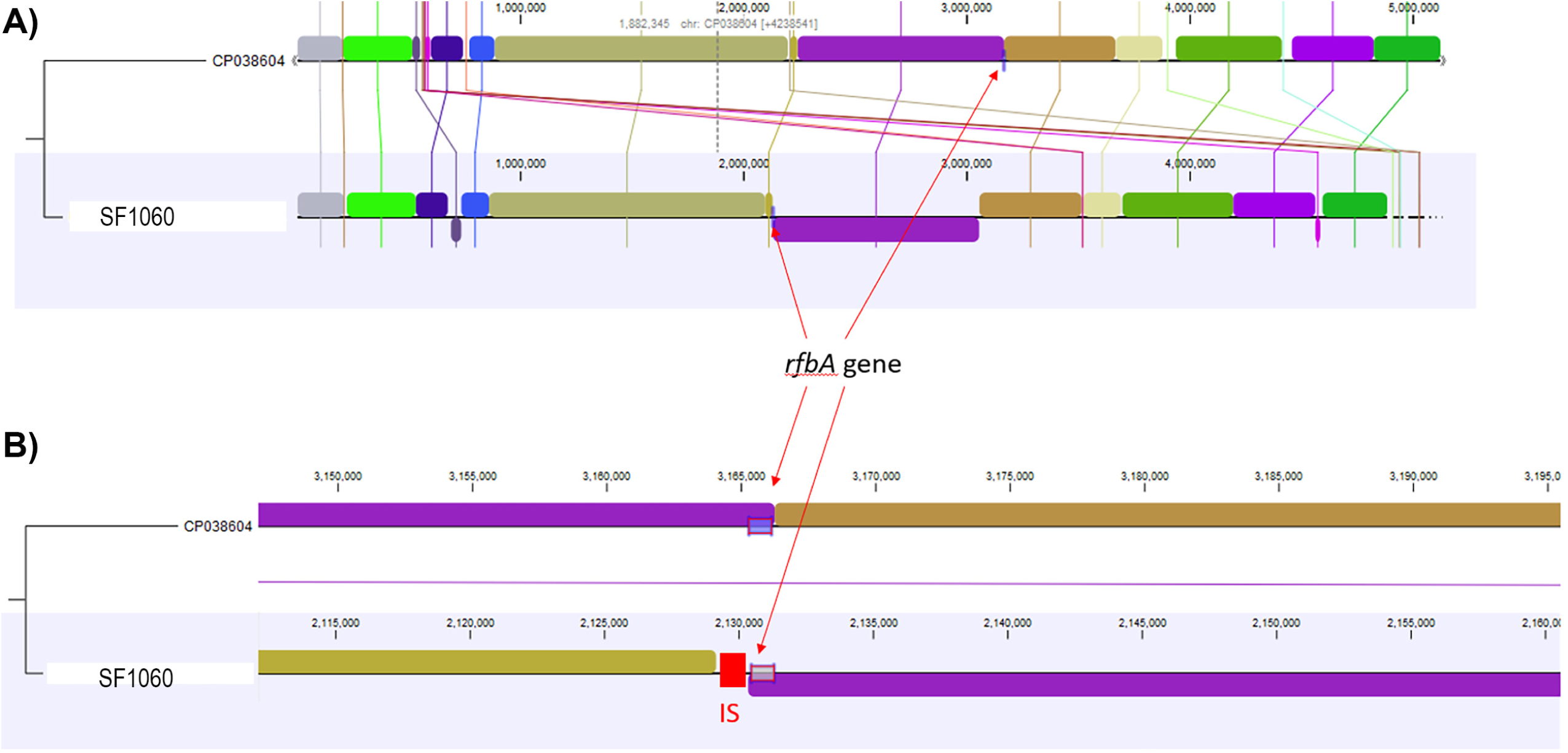
Chromosomal rearrangement at the *rfb* operon in *S. enterica* serovar Senftenberg strain SF1060 explains the loss of somatic antigen expression. Pairwise whole-genome alignment comparing SF1060 with *S.* Senftenberg strain GTA-FD-2016-MI-02533-2 (GenBank accession CP038604), which possesses an intact *rfb* operon and expresses normal O-antigens. The *rfb* gene cluster encodes enzymes required for the biosynthesis of O-antigen deoxysugars and lipopolysaccharide assembly. (A) Complete circular chromosome alignment showing a large-scale chromosomal inversion at the *rfb* locus in SF1060. Syntenic regions are connected by shaded bands, with inversions indicated by crossed bands. The rearrangement disrupts the *rfb* operon, preventing somatic antigen biosynthesis. (B) Detailed view of the *rfbD* gene region showing the mechanism of disruption. In the reference strain (top), the *rfb* operon (*rfbA, rfbB, rfbC, rfbD*) is intact. In SF1060 (bottom), an *IS5*-family transposase (*ISEc68* element, indicated by arrows and red colored boxes) has inserted at both ends of the inverted region, truncating the *rfbD* gene (encoding dTDP-4-dehydrorhamnose reductase) and separating it from the remaining *rfb* genes. This insertion sequence-mediated rearrangement renders the *rfb* operon non-functional, explaining the absence of detectable somatic antigens by conventional serotyping. The alignment was generated using CLC Genomics Workbench v20.

### Analysis of hydrogen sulfide production pathways

The *phs* operon of strain SF1060, responsible for reduction of thiosulfate to hydrogen sulfide, did not present premature stops but there were some SNPs differences when compared to the *phs* operon of *S.* Typhimurium LT2 (L32188.1) (Figure 3). SNPs are present in *phsA* and *phsC*. In the case of *phsA* there are missense mutations in position 420 (S>A), 421 (A>V) and in the last residue 758 (A>V). For *phsC* there are missense mutations in position 79 (L>F) and 156 (V>A).

**Figure 3.**
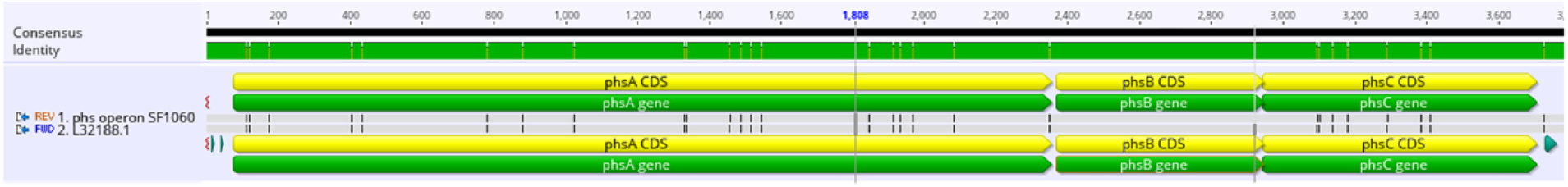
Alignment of the SF1060 *phs* operon with *S.* Typhimurium *phs* operon L32188.1. Comparative analysis of the *phs* operon in *S. enterica* serovar Senftenberg strain SF1060 and reference strain *S.* Typhimurium LT2. Nucleotide sequence alignment of the *phs* operon (thiosulfate reductase) from SF1060 compared to the reference *S.* Typhimurium LT2 phs operon (GenBank accession L32188.1). The *phs* operon comprises three genes (*phsA, phsB,* and *phsC*) responsible for the reduction of thiosulfate to hydrogen sulfide (H₂S). Despite the H₂S-negative phenotype of SF1060, no premature stop codons were identified in any of the *phs* genes. Missense mutations (indicated by arrows or highlighted positions) were detected in *phsA* at positions 420 (S>A), 421 (A>V), and 758 (A>V), and in *phsC* at positions 79 (L>F) and 156 (V>A). These amino acid substitutions may affect enzyme function, though their precise impact on H₂S production remains to be determined. The alignment was generated using Geneious Prime v2025.0.3.

The *ars* operon encodes proteins involved in the anaerobic reduction of sulphite, allowing the production of hydrogen sulphide (H₂S) from sulphite under oxygen-free conditions. In this operon there are only two missense mutations in position 208 and and 219 (H>R in both cases) of gene *arsA*.

### Phylogenetic relationship to regional and global *S.* Senftenberg strains

A wgMLST analysis showed that SF1060 clustered with another *S.* Senftenberg previously isolated in the region (also with the same chromosomal re-arrangement at the *rfb* operon) (Figure 4). The phylogenetic analysis included 24 *S.* Senftenberg genomes from diverse sources and geographic locations (Supplementary Table 1), including clinical strains, food samples, and wildlife sources from North America, South America, and Europe. *S.* Senftenberg is commonly isolated from seafood in northwestern Spain, where it accounts for up to 50% of the *S. enterica* strains identified. Although H₂S-negative variants have been previously reported, this new strain (SF1060), exhibits additional genetic modifications that complicate not only its detection but also its serotyping.

**Figure 4.**
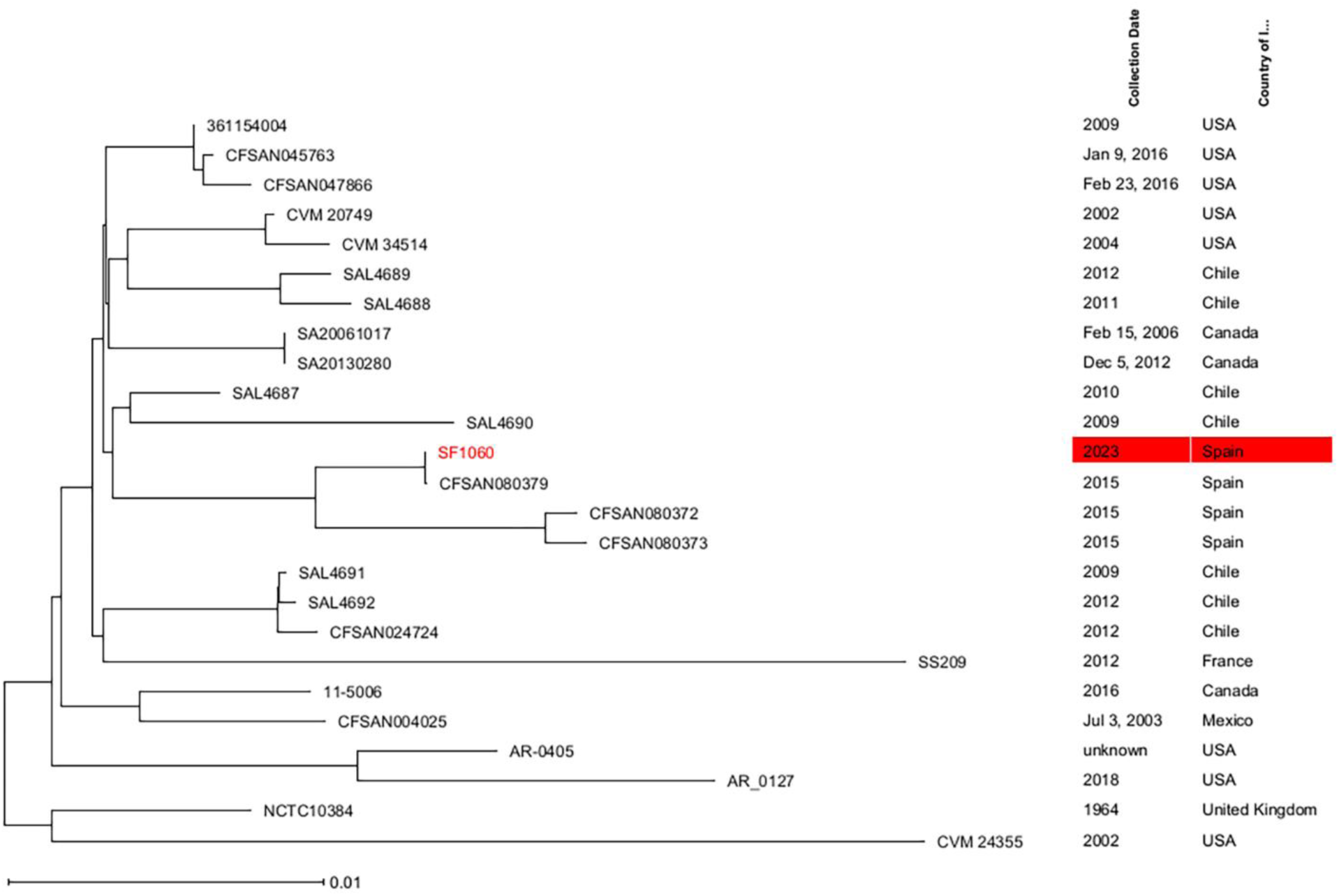
Phylogenetic relationship of *S. enterica* serovar Senftenberg strain SF1060 with global strains based on whole-genome multilocus sequence typing (wgMLST). Neighbor-joining (NJ) tree constructed using Nei’s DNA distance method based on allelic differences across 3,456 core genome coding sequences (CDSs) shared among SF1060 and 24 additional *S.* Senftenberg genomes retrieved from NCBI (Supplementary Table 1). The analysis included strains from diverse geographic locations (Spain, USA, Canada, Chile, France, United Kingdom) and sources (mussels, clinical samples, avian, food products) spanning 1964-2023. Strain SF1060 (highlighted in red) clustered with strain CFSAN080379 with no allelic differences detected by wgMLST, which was isolated from mussels in the same geographic region (Galicia, Spain) in 2015, indicating long-term persistence of this ST14 lineage in the regional marine aquaculture environment. The tree was constructed using Ridom SeqSphere+ v9.0.8 with pairwise ignoring of missing values. Branch lengths represent genetic distances based on the number of allelic differences in core genes.

## Discussion

This study identified and characterized an atypical *S.* Senftenberg strain (SF1060) from cooked mussels that presents dual challenges for conventional detection: loss of somatic antigen expression and absence of H₂S production. These traits, each with different genetic underpinnings, illustrate the complexity of *S. enterica* phenotypic variation and difficulties for correct isolation and identification using phenotypic methods. The loss of somatic antigen expression in SF1060 can be directly attributed to a chromosomal rearrangement within the *rfb* gene cluster, mediated by an *IS5*-family transposase (*ISEc68*). The *rfb* cluster encodes the enzymes involved in O-antigen biosynthesis, and variations within this cluster account for the diversity of somatic antigens observed across the *Salmonella* genus (16). This rearrangement truncated the *rfbD* gene, which encodes dTDP-4-dehydrorhamnose reductase, a critical enzyme in O-antigen biosynthesis, rendering the *rfbD* non-functional and explaining the serotyping failure. While other studies have reported serologically atypical *S.* Infantis isolated from poultry (17, 18); the underlying mechanism was not identified; the rearrangement found here presents a plausible explanation, underscoring the important role mobile genetic elements may play in creating antigenic diversity in *S. enterica* across various environments. Likewise, IS-mediated disruption of O-antigen biosynthesis has been documented in *E. coli* O157:H7, where IS629 insertion caused gene disruption rather than chromosomal rearrangement (19, 20).

In contrast, the mechanism for the H₂S-negative phenotype remains unclear. The reference media commonly used for *S. enterica* isolation is XLD agar, on which colonies typically appear black due to hydrogen sulfide (H₂S) production from thiosulfate. The H₂S reacts with iron salts in the medium to form the characteristic black precipitate, facilitating visual identification of presumptive *S. enterica* colonies (4). In recent decades, multiple strains of H₂S-negative *S. enterica* serotypes, including *S.* Infantis (21), *S.* Typhimurium (22), and *S.* Senftenberg (23) have been reported. Alterations in the pathways that reduce thiosulfate and sulfite to H₂S are believed to be responsible for the emergence of these strains. The *phs, cysJIH,* and *ars* operons are the main ones involved in this process: the *phs* operon mediates thiosulfate reduction, the *ars* operon reduces sulfite, and *cysJIH* is involved in the final steps of sulfite production (24). Among these, the *phs* operon is considered the most critical in the development of H_2_S-negative strains. H₂S-negative strains have been primarily associated with nonsense mutations in *phsA*, leading to a premature stop codon (21–23). However, in strain SF1060, no premature termination codons were identified in any of the *phs* operon genes; instead, three missense mutations were found in *phsA* (positions 420, 421, and 758) and two in *phsC* (positions 79 and 156). Similarly, the *ars* operon contained no nonsense mutations, though two missense mutations were observed in *arsA* (positions 208 and 219). No nonsense mutations were detected in the *cysJIH* operon either. These findings suggest that the absence of H₂S production in SF1060 is not attributable to the typical mechanisms involving premature stop codons in these operons. It is possible that the missense mutations identified affect protein folding, substrate binding, or catalytic efficiency, or alternatively, that regulatory mechanisms controlling operon expression are disrupted at the transcriptional or post-translational level, which will require future transcriptomic analysis to resolve.

The dual atypical phenotype of SF1060 exposes critical vulnerabilities in standard culture-based surveillance, which relies on colony morphology (e.g., black colonies on XLD) and serotyping for identification. Although molecular detection methods for *S. enterica*, primarily based on qPCR, have been developed in recent years, culture-dependent methods remain the gold standard for the detection of this pathogen (3, 4). These methods rely on a combination of non-selective and selective enrichment steps followed by plating on selective and differential agar media. The phenotypic characteristics of SF1060 expose critical vulnerabilities in these standard protocols. The absence of black colony formation on XLD agar could lead to false-negative results if laboratory personnel rely solely on typical colony morphology for presumptive identification. While SF1060 produced characteristic purple colonies on Brilliance™ *Salmonella* agar, demonstrating that chromogenic media can provide complementary detection capabilities, the lack of somatic antigen expression prevented conventional serotyping, which is essential for epidemiological tracking and source attribution. These limitations are not merely academic concerns; they have direct implications for outbreak investigations, where rapid and accurate strain characterization is critical for implementing control measures.

Molecular detection methods offer a robust solution to these challenges. The integration of rapid molecular screening techniques offers an effective strategy to overcome limitations inherent in culture-based diagnostics. Approaches such as quantitative PCR (qPCR) and loop-mediated isothermal amplification (LAMP) have proven particularly valuable in this context (25, 26), as they target conserved genetic sequences rather than phenotypic traits. In our study, qPCR successfully identified SF1060 as *S. enterica* despite its atypical characteristics, with a Cq value of 21 from the enriched mussel sample. However, these methods remain constrained by their limited ability to provide comprehensive strain characterization beyond detection. Hence, the implementation of whole-genome sequencing (WGS) workflows, coupled with user-friendly bioinformatics pipelines, presents a promising alternative for the detection, isolation, and detailed analysis of atypical strains (22, 27–31). WGS represents a more comprehensive approach, enabling simultaneous detection, serotyping, antimicrobial resistance profiling, and phylogenetic analysis from a single dataset.

In the case of SF1060, *in silico* serotyping using SeqSero2 correctly identified the strain as S. Senftenberg (1,3,19:g,s,t:-) and assigned it to ST14, information that was unattainable through conventional methods. Likewise, routine WGS enables the *in silico* identification of antimicrobial resistance and virulence determinants using databases such as ResFinder (14) and VFDB (32), respectively. The decreasing costs and increasing accessibility of long-read sequencing platforms, such as Oxford Nanopore Technologies, make WGS increasingly feasible for routine implementation in food safety laboratories (33–36). This technology also supports shotgun metagenomics approaches, enabling strain-level characterization of foodborne pathogens directly from outbreak samples (33, 35, 36). A multi-tiered surveillance approach is warranted: (i) molecular screening for rapid detection; (ii) multiple selective media to capture phenotypic variants; (iii) WGS-based characterization; and (iv) periodic protocol re-evaluation. This integrated strategy would enhance surveillance sensitivity and specificity while enabling effective risk management.

The persistence of *S.* Senftenberg ST14 in the mussel production environment of Galicia, Spain suggests it is an endemic pathogen adapted to this specific niche. It has been the predominant serotype isolated from the region’s shellfish and processing facilities for over two decades (37, 38). Whole-genome multi-locus sequence typing (wgMLST) analysis confirms this long-term persistence in the mussel production environment and its ongoing potential to contaminate shellfish, showing that recently isolated strain SF1060 is closely related to another ST14 strain (CFSAN080379) from 2015, with both sharing an identical chromosomal rearrangement at the *rfb* operon. The unique selective pressures of the marine environment, such as high salinity, characteristic of mussel production systems, may favor this lineage (39). Many of these regional strains exhibit a rough morphotype associated with an altered lipopolysaccharide structure, which enhances biofilm formation and environmental survival (37, 38). The presence of other atypical *S. enterica* strains, including some with resistance to last-resort antibiotics, has also been reported in this area (7).

Given that Galicia’s mussel aquaculture accounts for over 90% of Spanish production, the presence of this pathogen poses a significant public health and economic risk (5). *S.* Senftenberg is a notable cause of human salmonellosis in the European Union, with 101 confirmed cases in 2023 (1), and its detection in cooked mussels indicates either post-processing contamination or inadequate thermal treatment. The atypical phenotypes of these persistent strains may lead to systematic underreporting, complicating control efforts. Therefore, enhanced surveillance is essential, requiring an integrated approach that combines targeted monitoring of environmental reservoirs and processing facility biofilms with genomic surveillance to track persistent lineages. Such measures are critical in coastal regions where aquaculture, tourism, and urban development create complex interfaces between human activities and marine ecosystems.

The *IS5*-family transposase-mediated chromosomal rearrangement in SF1060 illustrates how mobile genetic elements generate phenotypic diversity within *S. enterica* populations. Such rearrangements may affect not only serological classification but also host-pathogen interactions and environmental fitness. However, culture-based surveillance methods systematically exclude strains lacking typical phenotypic markers, potentially leading to substantial underreporting of atypical variants. Culture-independent methods, such as shotgun metagenomics applied directly to food samples (36, 40–44), could reveal the true extent of this hidden diversity and provide more comprehensive pathogen surveillance.

Future research should elucidate the molecular basis of the H₂S-negative phenotype through transcriptomic and proteomic analyses of the observed missense mutations. Additionally, experimental evolution studies could assess whether *rfb* rearrangements confer fitness advantages under marine aquaculture conditions, such as high salinity or biofilm formation. Development of predictive algorithms to infer phenotypes from genomic sequences would enable rapid assessment of potential detection challenges for emerging variants. Integration of genomic data into international surveillance platforms such as NCBI Pathogen Detection (https://www.ncbi.nlm.nih.gov/pathogens), enables real-time comparison of isolates against global databases, facilitating identification of related strains, outbreak detection, and monitoring of lineage spread. By linking strains from diverse sources (including food, clinical cases, wildlife, and environmental samples) these platforms support a One Health approach (27, 45–49) to understanding *S. enterica* epidemiology, tracing transmission pathways, and identifying critical control points across the human-animal-environment interface.

## Conclusions

The loss of key phenotypic traits, such as the ability to metabolize thiosulfate to hydrogen sulfide, and chromosomal rearrangements leading to the absence of somatic antigen expression present significant challenges for the isolation and characterization of *S. enterica* using conventional phenotypic methods. These limitations underscore the need to integrate whole-genome sequencing into routine surveillance and monitoring programs. The adoption of WGS not only enhances the detection and precise characterization of atypical strains but also strengthens outbreak investigations and contributes to a more comprehensive understanding of *S. enterica* evolution and persistence in the food chain. Such an integrated approach is essential to reinforce surveillance systems, improve outbreak response capacity, and ensure the continued protection of public health. Furthermore, the endemic persistence of atypical *S.* Senftenberg variants in marine aquaculture environments highlights the need for One Health surveillance frameworks that integrate genomic data from food, environmental, and clinical sources to effectively track pathogen evolution and transmission across ecological niches.

## Ethical approval and consent to participate

N/A

## Consent for publication

N/A

## Author contributions

Conceptualization: A. L., A. L-L., A. G-M., N. G-E., Data curation: A. L., A. G-M., N. G-E., Formal analysis: A. L., A. G-M., N. G-E., N. G-E, Investigation: A. G-M., N. G-E., Methodology: A. L., A. L-L., A. G-M., N. G-E, Project administration: A. G-M., N. G-E., Resources: A. G-M., N. G-E., Supervision: A. G-M., N. G-E., Validation: A. L., A. L-L., A. G-M., N. G-E., Visualization: A. L., A. L-L., A. G-M., N. G-E., Writing – original draft: A. L., A. L-L., A. G-M., N. G-E., Writing – review and editing: A. L., A. L-L., A. G-M., N. G-E., Funding acquisition: A. G-M.

## Acknowledgements

We thank the staff at SGS Seafood Laboratory for sample collection and the Veterinary Health Surveillance Center (VISAVET) of the Complutense University of Madrid for serotyping analysis. We acknowledge Biomarker Technologies (BMKGENE GmbH, Münster, Germany) for Illumina sequencing services.

## Data availability

The complete genome sequence of *S. enterica* serovar Senftenberg strain SF1060 has been deposited in GenBank under BioProject accession number PRJNA1365827, BioSample accession number SAMN53295204. Raw sequencing data are available under SRA accession numbers SRR36084183 (Illumina) and SRR36084182 (Oxford Nanopore Technologies). The assembled genome is available under accession number ACBLBR000000000. All data supporting the findings of this study are available within the article and its supplementary materials.

